# Distinct endocannabinoids specifically signal to astrocytes and neurons

**DOI:** 10.1101/2023.06.13.544877

**Authors:** Jose Antonio Noriega-Prieto, Rafael Falcón-Moya, Abel Eraso-Pichot, Unai B Fundazuri, Pavan Guttipatti, Lindsey Belisle, Antonio Rodríguez-Moreno, Mario van der Stelt, Joseph Cheer, Giovanni Marsicano, Paulo Kofuji, Alfonso Araque

## Abstract

The endocannabinoid system is an essential intercellular signaling mechanism with a decisive role in many physiological functions of the brain. Endocannabinoids (eCBs), directly acting on presynaptic neuronal CB1 receptors (CB1Rs), can inhibit neurotransmitter release. In addition, they can potentiate adjacent synapses, inducing lateral regulation of synaptic transmission through astrocyte CB1Rs. In contrast to most, if not all, neurotransmitter systems, the eCB system involves two distinct ligands, Anandamide and 2-Arachidonoylglycerol (AEA and 2AG), and a single receptor (CB1R). The physiological meaning of this particularity remains unknown. Here we show that different eCBs are signaling both astrocytes and neurons, inducing distinct and contrasting synaptic regulation. Combining two-photon with a pharmacological and optogenetic approaches and transgenic mice for the synthesis enzyme of both eCBs, we have found that the absence of 2-AG synthesis abolished the inhibitory effect, which was mediated exclusively by neuronal mechanisms. However, the absence of AEA synthesis prevents the lateral potentiation mediated by astrocyte calcium mobilization. Together this indicates that 2-AG signals to neurons, decreasing neurotransmitter release, while AEA signals to astrocytes and induces lateral potentiation. Additionally, AEA synthesis is required for the synaptic potentiation induced by spike-timing-dependent plasticity, as well as astrocyte CB1R, indicating that distinct eCBs-signaling influences neuronal plasticity. We conclude that 2-AG and AEA induce distinct and contrasting synaptic regulation through CB1R in different cell types.

## Introduction

The endocannabinoid system is a significant intercellular signaling system with crucial function in the regulation of synaptic plasticity and control of behavior throughout the central nervous system (Ahn et al., 2008; Alger & Kim, 2011; Castillo et al., 2012; Di Marzo, 2009; Marcus et al., 2020; Martin-Fernandez et al., 2017a; Ohno-Shosaku & Kano, 2014; Piomelli, 2003; Robin et al., 2018; Serrat et al., 2022; Skupio et al., 2023). The increase in postsynaptic activity induces the mobilization of intracellular postsynaptic calcium causing the synthesis of 2-Arachidonoylglycerol (2-AG) by diacylglycerol lipase (DGL) and anandamide (AEA) by N-acetylphosphatidylethanolamine-hydrolysing phospholipase D (NAPE-PLD), the most prominent endocannabinoids (eCBs) in the central nervous system (CNS) (Noriega-Prieto et al., 2023). The particular retrograde action of eCBs is exerted through presynaptic cannabinoid receptor type 1 (CB1R) expressed by neurons resulting in a decrease in release of neurotransmitter (Castillo et al., 2012). Additionally, eCBs can regulate synaptic function through astroglial CB1R inducing lateral regulation (Baraibar et al., 2022; Covelo & Araque, 2018; Martín et al., 2015; Martin-Fernandez et al., 2017a; Navarrete & Araque, 2010a). Ultimately, eCB signaling concludes by the reuptake of the eCBs and the enzymatic degradation of 2-AG by monoacylglycerol lipase (MGL) (Dinh et al., 2002) and AEA by fatty acid amide hydrolase (FAAH) (Cravatt et al., 1996; Hillard et al., 1995; McKinney & Cravatt, 2005).

Astrocytes are active cells that participate in the regulation of synaptic physiology and information processing. They sense neurotransmitters expressing a plethora of G protein-coupled receptors in the membrane, which induce intracellular calcium variations, producing, in turn, the release of gliotransmitters such as glutamate, d-serine and ATP (Andrade-Talavera et al., 2016; Covelo & Araque, 2018; Martin-Fernandez et al., 2017a; Navarrete & Araque, 2010a; Robin et al., 2018). The astrocytic release of gliotransmitters activates neuronal receptors, regulating synaptic transmission and plasticity. Astrocytes respond to eCBs through activation of PTX-insensitive G protein CB1Rs expressed in their membrane, which is a prominent modulatory mechanism (Navarrete & Araque, 2008, 2010a). It has been described that eCB-induced astrocyte activation produces lateral regulation of synaptic plasticity in different brain areas with behavioral repercussions (Navarrete and araque 2010, martin et al., 2015, Martin-Ferandez et al.(Baraibar et al., 2022; Covelo & Araque, 2018; Martín et al., 2015; Martin-Fernandez et al., 2017a; Navarrete & Araque, 2010a), which define a differential, but not exclusive, astrocytic vs. neuronal mechanism induced by eCBs.

In contrast to most, if not all, neurotransmitter systems that involve a single ligand and numerous receptor types, the eCB system involves two distinct ligands (AEA and 2AG) and a single receptor (CB1Rs). The physiological meaning of this particularity remains unknown. Despite the rising evidence demonstrating that eCB can signal either astrocytes or neurons (Noriega-Prieto et al., 2023), the cellular mechanism is still unknown. We investigated the differential eCBs signaling to astrocytes and neurons through neuronal and astrocytic CB1R at hippocampal CA1 pyramidal neurons. Using electrophysiological recordings, two-photon calcium imaging, pharmacological and optogenetic approaches and transgenic mice targeting the synthesis enzyme of both eCBs, 2-AG and AEA, we performed paired recordings from CA1 pyramidal neurons, monitored astrocyte Ca2+ levels, and stimulated Schaffer collaterals using the minimal stimulation technique that activates single or very few presynaptic fibers (Dobrunz & Stevens, 1997; Isaac et al., 1996; Perea & Araque, 2007; Raastad, 1995). We have found that AEA specifically signals to astrocytes and 2-AG to neurons leading to distinct and contrasting synaptic regulation in single hippocampal synapses. In addition, the distinct astrocyte-driven signaling impacts synaptic plasticity induced by spike-timing dependent plasticity (STDP) which further supports the idea that astrocyte function influences synaptic activity through eCB signaling.

## Materials and Methods

### Ethics Statement

All animal care and sacrifice procedures were approved by the University of Minnesota Institutional Animal Care and Use Committee (IACUC) with compliance to the National Institutes of Health guidelines for the care and use of laboratory animals. Mice were housed under 12/12-h light/dark cycle and up to five animals per cage. The following animals (males and females) were used for the present study C57BL/6J, NAPE-PLD−/− and DAGLα−/− (generously donated by Dr. Cravatt), CB1R flox/flox and IP3R2−/−. Adult (≥6 weeks) mice were used.

## Method Details

### Slice Preparation

Animals were decapitated, and the brains were submerged in cold (4°C) cutting solution containing the following (in mM): 189.0 sucrose, 10.0 glucose, 26.0 NaHCO_3,_ 3.0 KCl, 5.0 Mg_2_SO_4_, 0.1 CaCl_2_, and 1.25 NaH_2_PO_4_.2H_2_O. Coronal slices (350 µm thick) were cut using a vibratome (model VT 1200S, Leica), and incubated (>1 h, at 25–27°C) in artificial CSF (ACSF) containing the following (in mM): NaCl 124, KCl 2.69, KH_2_PO_4_ 1.25, MgSO_4_ 2, NaHCO_3_ 26, CaCl_2_ 2, and glucose 10, and oxygenated with 95% O_2_ / 5% CO_2_ (pH = 7.3-7.4). Slices were placed in an immersion recording chamber and superfused (3 mL/min) with oxygenated ACSF at 32-34 °C and visualized with an Olympus BX50WI microscope (Olympus Optical, Japan), a custom made confocal microscope (Thorlabs) or a two-photon microscope (model DM6000 CFS upright multiphoton microscope with TCS SP5 MP Laser, Leica).

### Electrophysiology

The whole-cell patch clamp technique was used to make electrophysiological recordings of CA1 neurons of the hippocampus. When filled with an internal solution containing (in mM): KGluconate 135, KCl 10, HEPES 10, MgCl 1, ATP-Na2 2 (pH = 7.3) (pH = 7.3), patch electrodes resistance ranged between 3-10 MΩ. The membrane potential was held at −70 mV. Series and input resistances were monitored throughout the experiment using a −5 mV pulse. Signals were recorded with PC-ONE amplifiers (Dagan Instruments, MN, US) and fed to a Pentium-based PC through a DigiData 1440A interface board. Signals were filtered at 1 KHz and acquired at 10 KHz sampling rate. The pCLAMP 11 (Molecular Devices) software was used for stimulus generation, data display, acquisition and storage.

### Synaptic stimulation and drug application

Synaptic currents were evoked using bipolar theta capillaries filled with ACSF placed in the brain region of study (CA1 of the hippocampus). Paired pulses (1 ms duration with 50 ms interval) were continuously delivered at 0.33 Hz using a stimulator S-910 (Dagan Instruments) through an isolation unit. Excitatory post-synaptic currents (EPSCs) were isolated using picrotoxin (50 μM) and CGP5462 (1 μM) to block GABA_A_R and GABA_B_R, respectively. The parameters analyzed were mean amplitude of EPSC response and paired pulse ratio (PPR = 2^nd^ EPSC/1^st^ EPSC). EPSC amplitudes were grouped in 10 s time bins, baseline mean EPSC amplitude was obtained by averaging mean values obtained within 3 min of baseline recordings and mean EPSC amplitudes were normalized to baseline. Stimulus effects were statistically tested comparing the normalized EPSCs recorded 30 s before and after the stimulus to assess changes in EPSC amplitude and PPR.

### Ca^2+^ imaging

Cytoplasmic Ca^2+^ levels in astrocytes in the CA1 of the hippocampus were monitored using two photon and confocal microscopy. Cells were illuminated during 100-200 ms with a laser at 488 nm (Thorlabs) and images were acquired every 1-2 s. The laser and the CCD camera were controlled and synchronized by the MetaMorph software (Molecular devices). Two photon or Confocal imaging utilized an Olympus BX61Wl microscope (Olympus Optical, Japan) controlled by Leica SP5 multi-photon microscope (Leica Microsystems, USA) or BX51Wl controlled by the ThorImageLS software. For control and pharmacology experiments Ca^2+^ was monitored before and after optostimulation of CA1 pyramidal neurons, using the genetically encoded Ca^2+^ indicator dye GCaMP6 under the GfaABC1D promoter to specifically target astrocytes. For optical stimulation of CA1 pyramidal neurons, a fiber optic connected to an LED (595 nm) was placed over the CA1 and a 5 s duration light was applied, using the channelrhodopsin ChrimsonR under the neuronal promoter synapsin. Note that Ca^2+^ experiments were performed in the presence of a cocktail of neurotransmitter receptor antagonists containing: CNQX (20 μM), AP5 (50 μM), MPEP (50 μM), LY367385 (100 μM), picrotoxin (50 μM), CGP5462 (1 μM), atropine (50 μM), CPT (10 μM), suramin (100 μM) and flupenthixol (30 μM).

Videos were obtained at 512_×_512 resolution with a sampling interval of 1-2 s. A custom MATLAB program (Calsee: https://www.araquelab.com/code/) was used to quantify fluorescence level measurements in astrocytes. Ca^2+^ variations recorded at the soma and processes of the cells were estimated as changes of the fluorescence signal over baseline (ΔF/F_0_), and cells were considered to show a Ca^2+^ event when the ΔF/F_0_ increase was at least two times the standard deviation of the baseline. The astrocyte Ca^2+^ signal was quantified from the Ca^2+^ event probability, which was calculated from the number of Ca^2+^ elevations grouped in 10 s bins recorded from 2-50 astrocytes per field of view. For each astrocyte analyzed, values of 0 and 1 were assigned for bins showing either no response or a Ca^2+^ event, respectively, and the Ca^2+^ event probability was obtained by dividing the number of astrocytes showing an event at each time bin by the total number of monitored astrocytes. To examine the difference in Ca^2+^ event probability in distinct conditions, the basal Ca^2+^ event probability (10 s before a stimulus) was averaged and compared to the average Ca^2+^ event probability (10 s after a stimulus).

### Stereotaxic Surgery

Adult mice were anesthetized with a ketamine (100 mg/kg)/ xylazine (10mg/kg) cocktail. Viral vectors (0.5 μl-1 μl) were injected bilaterally using a Hamilton syringe attached to a 29-gauge needle at a rate of 0.1 μl/min. The viral constructs AAV5-GfaABC1D-cytoGCaMP6f (Addgene) was targeted to CA1 astrocytes (anterior-posterior [AP]: – 2.3 mm; medial-lateral [ML]: +/− 1.5 mm; dorsal-ventral [DV]: −1.2 mm) of C57BL/6J, NAPE-PLD−/−, DAGLα−/− and CB1R flox/flox. The AAV8-GFAP-mCherry-Cre or AAV8-GFAP-mCherry (UMN vector core) were targeted to CB1R flox/flox astrocytes. The AAV5-Syn-ChrimsonR-tdT (Addgene) was also targeted to CA1 neurons in the hippocampus of all strains. Mice were used ≥ 2 weeks after stereotaxic surgeries.

### Drugs

4-[3-[2-(Trifluoromethyl)-9*H*-thioxanthen-9-ylidene]propyl]-1-piperazineethanol dihydrochloride (flupenthixol dihydrochloride) from abcam. [*S*-(*R*^∗^,*R*^∗^)]-[3-[[1-(3,4-Dichlorophenyl)ethyl]amino]-2-hydroxypropyl](cyclohexylmethyl) phosphinic acid (CGP 54626 hydrochloride) and (*S*)-(+)-α-Amino-4-carboxy-2-methylbenzeneacetic acid (LY367385) from Tocris. 8,8’-[Carbonyl*bis*[imino-3,1-phenylenecarbonylimino(4-methyl-3,1-phenylene)carbonylimino]]*bis*-1,3,5-naphthalenetrisulfonic acid hexasodium salt (suramin hexasodium salt) from Thermo Fisher Scientific. *N*-(Piperidin-1-yl)-5-(4-iodophenyl)-1-(2,4-dichlorophenyl)-4-methyl-1*H*-pyrazole-3-carboxamide (AM 251), D-(-)-2-Amino-5-phosphonopentanoic acid (D-AP5), 6-Cyano-7-nitroquinoxaline-2,3-dione disodium (CNQX disodium salt), and 2-Methyl-6-(phenylethynyl)pyridine hydrochloride (MPEP hydrochloride) were purchased from Cayman. Picrotoxin from Sigma-Aldrich.

### Quantification and Statistical Analysis

Data are expressed as mean ± standard error of the mean (SEM). For electrophysiology comparisons number of neurons was used as the sample size; for *in vitro* Ca^2+^ signal comparisons the number of slices was used as the sample size. At least 3 mice per experimental group were used. Data normality was tested using a Shapiro-Wilk test. Results were compared using a two-tailed Student’s t test or ANOVA (α = 0.05). One-way ANOVA with a Holm-Sidak’s multiple comparisons test post hoc was used for normal distributed data and Kruskal-Wallis One-Way ANOVA with Dunn’s method post hoc was used for non-normal distributed data. Statistical differences were established with p < 0.05 (^∗^), p < 0.01 (^∗∗^) and p < 0.001 (^∗∗∗^).

## Results

### 2-AG but not AEA Transiently Decreases Neurotransmitter Release at Hippocampal CA3-CA1 Synapses

The endocannabinoids (eCBs) have been described to signal to both astrocytes and neurons inducing phenomena called “endocannabinoid-mediated the synaptic potentiation” (eSP) and depolarization-induced suppression of excitation (DSE), respectively, through the CB1R activation (Covelo & Araque, 2018; Navarrete & Araque, 2010a). While the role of 2-AG in inducing DSE in various regions of the central nervous system (CNS) has been well-documented, the modulation of synaptic transmission by anandamide (AEA) is still not fully understood.

We conducted pair recordings in CA1 pyramidal neurons of the hippocampus (Fig. 1A) to assess synaptic efficacy of EPSCs induced by minimal stimulation in both neurons. After establishing a stable baseline of EPSCs, we depolarized (ND) one neuron to 0 mV for 5 seconds to elicit the release of endocannabinoids (eCBs). This depolarization (ND) resulted in depolarization-induced suppression of excitation (DSE) in the stimulated neuron or homoneuron (from 98.93 ± 2.82 to 66.73 ± 5.61% of peak amplitude, n=26, ***p<0.001; Fig. 1B and C). To confirm that DSE was mediated by CB1R activation, we blocked the CB1R receptor using the antagonist AM-251, which abolished the DSE effect (from 100.70 ± 2.82 to 100.61 ± 5.52% of peak amplitude, n=7, p=0.99; Fig. 1E; AM-251). To investigate the involvement of astrocytes in DSE, we selectively deleted the CB1R receptor in hippocampal astrocytes (GFAP-CB1R-/-mice) using a Cre/lox approach via viral injection. Interestingly, DSE was still induced both in GFAP-CB1R+/+ (from 107.8 ± 3.88 to 67.97 ± 4.15% of peak amplitude, n=5, ***p<0.001; Fig. 1E; GFAP-CB1R+/+) and GFAP-CB1R-/-(from 99.34 ± 0.65 to 60.29 ± 4.94% of peak amplitude, n=5, ***p<0.001; Fig. 1E; GFAP-CB1R-/-) mice. These findings suggest that astrocytic CB1R is not involved in mediating DSE.

**Figure 1.**
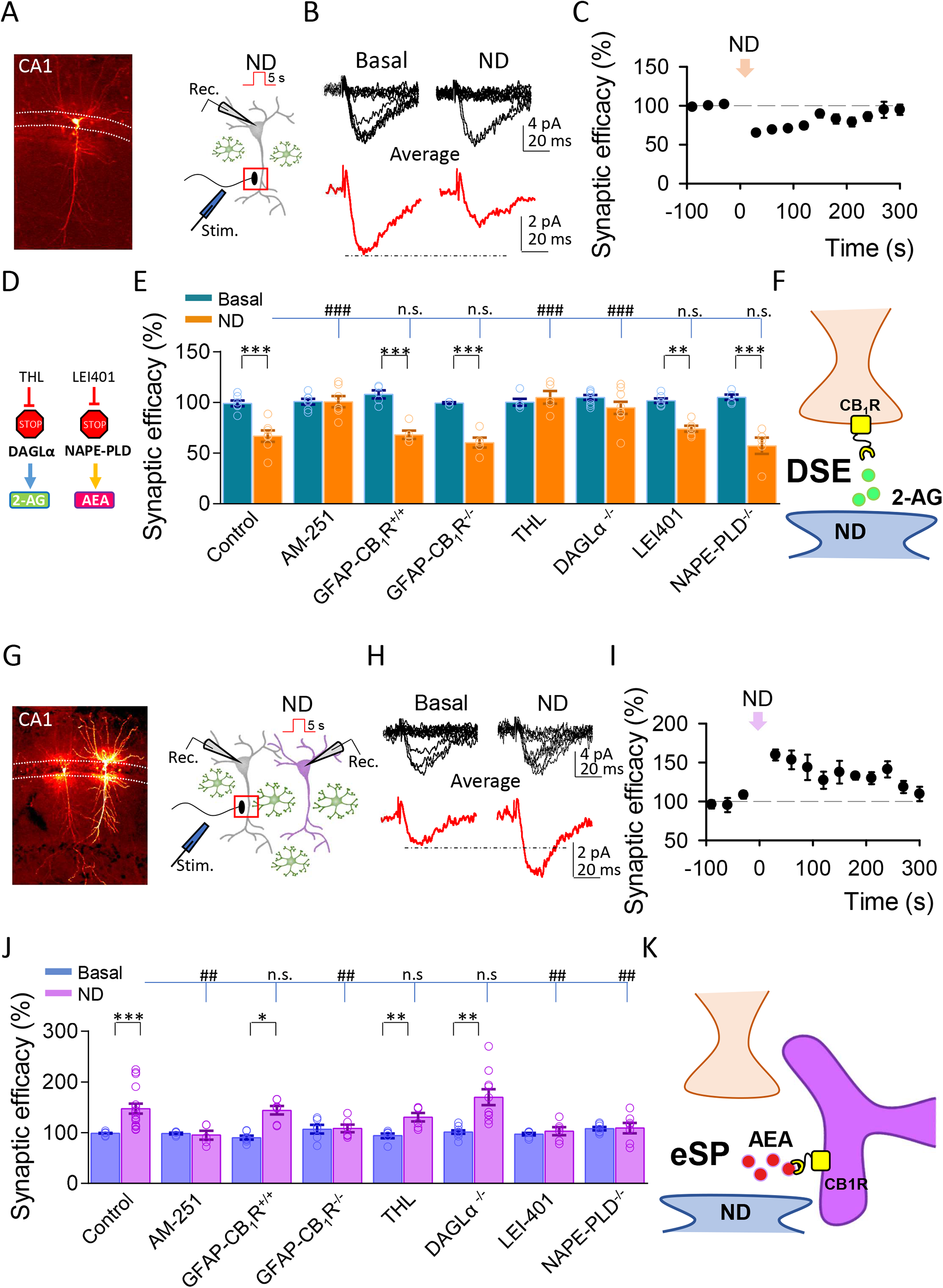
2-AG signals to neurons and depresses excitatory synaptic transmission and AEA signals to astrocytes and enhances excitatory synaptic transmission in CA1 synapses. **(a)** left. Biocytin loading CA1 pyramidal neuron image. Right, schematic representation of the experimental design. (**b**) EPSCs evoked by minimal stimulation showing EPSC amplitudes and failures of synaptic transmission (20 consecutive stimuli) (top traces) and averaged EPSCs (n = 20 stimuli, including successes and failures; bottom red traces) before (basal) and after homoneuron ND. (Scale bars: 4 pA, 20 ms for EPSCs and 2 pA, 20 ms for average traces). (**c**) Time course of synaptic efficacy before and after ND in control condition of the homoneuron. (**d**) Schematic representation of synthesis profile of 2-AG and AEA. (**e**) Relative changes from control basal values of synaptic efficacy before and after homoneuron ND in control conditions, AM251, GFAP-CB1R^+/+^, GFAP-CB1R^-/-^, THL, DAGLα^-/-^, LEI401 and NAPE-PLD^-/-^. (**f**) Schematic drawing representing that after neuronal depolarization (ND), 2-AG release signals to neurons through presynaptic CB1R and induces DSE. (**g**) left. Biocytin loading CA1 pyramidal neurons image. Right, schematic representation of the experimental design of neuronal pair recording. (**h**) EPSCs evoked by minimal stimulation showing EPSC amplitudes and failures of synaptic transmission (20 consecutive stimuli) (top traces) and averaged EPSCs (n = 20 stimuli, including successes and failures; bottom red traces) before (basal) and after ND of the heteroneuron. (Scale bars: 4 pA, 20 ms for EPSCs and 2 pA, 20 ms for average traces). (**i**) Time course of synaptic efficacy before and after ND in control condition of the heteroneuron. (**j**) Relative changes from control basal values of synaptic efficacy before and after heteroneuron ND in control conditions, AM251, GFAP-CB1R^+/+^, GFAP-CB1R^-/-^, THL, DAGLα^-/-^, LEI401 and NAPE-PLD^-/-^. (**k**) Schematic drawing representing that AEA signals to astrocytes and induces e-SP. AEA released activates astroglial CB1Rs, decreasing neurotransmitter release.

Previous studies have demonstrated that 2-AG is responsible for inducing depolarization-induced suppression of excitation (DSE) (Ohno-Shosaku & Kano, 2014). 2-AG is synthesized from diacylglycerol (DAG), with DAGLα being the isoform involved in mediating 2-AG-mediated synaptic transmission in the adult brain (Murataeva et al., 2014). In our study, we aimed to investigate the specific endocannabinoid (eCB) involved in DSE. By using the antagonist of DAGLα (Fig. 1D, DAGLα) which is the synthesis enzyme for 2-AG, known as THL (Fig. 1D; THL), we were able to prevent the occurrence of DSE (from 100.05 ± 3.44 to 104.83 ± 6.29% of peak amplitude, n=5, p=0.63; Fig. 1E; THL). Additionally, we observed the absence of DSE in DAGLα-/-mice, which further confirms the role of 2-AG as the essential eCB for DSE (from 104.77 ± 2.23 to 94.62 ± 5.87% of peak amplitude, n=9, p=0.11; Fig. 1E; DAGLα-/-). Interestingly, the presence of the NAPE-PLD blocker (Fig. 1D, NAPE-PLD), which is the synthesis enzyme of AEA, namely LEI 401 (Fig. 1D, LEI 401), did not affect DSE (from 101.69 ± 2.32 to 73.92 ± 2.94% of peak amplitude, n=6, **p<0.01; Fig. 1E; LEI 401), and this observation was further confirmed in NAPE-PLD-/-mice (from 105.01 ± 2.58 to 57.14 ± 8.06% of peak amplitude, n=5, **p<0.01; Fig. 1E; NAPE-PLD-/-). Additionally, the transient reduction of synaptic transmission exhibited presynaptic expression, as indicated by the change in paired-pulse ratio (PPR) (from 102.25 ± 2.98 to 132.63 ± 7.96%, n=26, **p<0.01; Supplementary Fig. 1A). These findings suggest that the postsynaptic induction of 2-AG release is exclusively due to neuronal signaling, leading to DSE (Fig. F). Furthermore, it indicates that AEA does not participate in DSE in this specific synapse.

### AEA but not 2-AG Transiently Potentiates Neurotransmitter Release through Hippocampal Astrocytes

Our previous findings in 2010 established the fundamental understanding of endocannabinoid-induced synaptic potentiation (eSP) mediated through astrocyte CB1R in the hippocampus (Navarrete & Araque, 2010a). This phenomenon, known as eSP, has been subsequently observed in various brain regions, including the basal ganglia, amygdala, and cortex, indicating it as a widespread mechanism of synaptic regulation (Baraibar et al., 2022; Martín et al., 2015; Martin-Fernandez et al., 2017a). However, the specific endocannabinoid responsible for inducing this lateral regulation of synaptic transmission remains unknown.

To address this question, we conducted pair recordings of CA1 pyramidal neurons in the hippocampus and examined the synaptic efficacy of the adjacent heteroneuron (Fig. 1G). Following a baseline measurement of EPSCs, depolarization (ND) resulted in an increase in synaptic efficacy of the heteroneuron (from 98.61 ± 0.69 to 147.32 ± 9.80 % of peak amplitude, n=25, ***p<0.001; Fig. 1H and I). This increase was blocked by the CB1R antagonist, AM-251 (from 98.01 ± 1.74 to 94.92 ± 8.75 % of peak amplitude, n=5, p>0.99; Fig. 1J; AM-251), indicating that e-SP was induced through CB1R activation. However, while e-SP persisted in GFAP-CB1R+/+ mice (from 90.03 ± 3.94 to 143.97 ± 8.29 % of peak amplitude, n=7, *p<0.05; Fig. 1J; GFAP-CB1R+/+), it was absent in mice with specific removal of CB1R in hippocampal astrocytes (from 106.65 ± 8.65 to 108.16 ± 7.62 % of peak amplitude, n=6, p>0.99; Fig. 1J; GFAP-CB1R-/-).

Furthermore, we observed that e-SP was still induced in the presence of THL, a blocker of the synthesis enzyme for 2-AG (from 93.76 ± 4.53 to 140.12 ± 7.71 % of peak amplitude, n=6, ***p<0.001; Fig. 1J; THL), and in DAGLα-/-mice, which lack the isoform involved in 2-AG synthesis (from 101.01 ± 3.24 to 179.68 ± 15.63 % of peak amplitude, n=10, **p<0.01; Fig. 1J; DAGLα-/-). Interestingly, the antagonist of NAPE-PLD, LEI-401, abolished e-SP (from 96.77 ± 2.68 to 102.89 ± 7.99 % of peak amplitude, n=6, p=0.55; Fig. 1J; LEI-401), as well as in NAPE-PLD-/-mice, which lack the synthesis enzyme of AEA (from 107.89 ± 3.26 to 109.08 ± 10.40 % of peak amplitude, n=7, p=0.91; Fig. 1J; NAPE-PLD-/-). These findings suggest that the postsynaptic induction of AEA release signals to astrocytes and mediates e-SP through astrocyte activation. Additionally, the change in the paired-pulse ratio (PPR) indicates a presynaptic origin of the expression of e-SP (from 102.50 ± 1.90 to 85.06 ± 4.34 % of peak amplitude, n=25, **p<0.01; Supplementary Fig. 1B).

In summary, our findings suggest that when the postsynaptic neuron is stimulated, it leads to the release of AEA, which then activates astrocytes through the CB1R. This activation of astrocytes results in a temporary increase in the release of glutamate from neurons.

### Optogenetic Stimulation of Pyramidal Neurons Induces Astrocyte Ca^2+^ Mobilization

Previous studies have demonstrated that astrocytes have a relatively low expression of CB1 receptors (CB1Rs) (Covelo et al., 2021). However, the functionality of astrocytic CB1Rs is crucial for various physiological changes in synaptic transmission, synaptic plasticity, and behavior (Baraibar et al., 2022; Covelo & Araque, 2018; Gómez-Gonzalo et al., 2015; Martin-Fernandez et al., 2017a; Navarrete & Araque, 2010a; Robin et al., 2018; Serrat et al., 2022; Skupio et al., 2023). Activation of CB1Rs in astrocytes and the subsequent mobilization of intracellular calcium play a vital role in modulating synaptic transmission and plasticity (Covelo & Araque, 2018; Gómez-Gonzalo et al., 2015; Martin-Fernandez et al., 2017b; Navarrete & Araque, 2010b). In our experiment, we employed specific tools to investigate the role of astrocytes in calcium activity and synaptic potentiation. We used the genetically encoded calcium sensor GCaMP6f under the astrocyte-specific promoter GfaABC1D to monitor calcium activity in astrocytes. Additionally, we utilized ChrimsonR, a light-gated cation channel, as an optogenetic tool to depolarize neurons simultaneously monitoring several synapsis, and induce synaptic potentiation (Fig. 2A and B) (Klapoetke et al., 2014). Through this setup, we observed an increase in synaptic potentiation from 109.46 ± 3.65 to 168.80 ± 10.35% of amplitude (n=3, *p<0.05; Supplementary. Fig. 2). To ensure that the observed effects were specifically due to astrocyte calcium activity, we employed a cocktail of neurotransmitter receptor antagonists (CNQX [20 μM], AP5 [50 μM], MPEP [50 μM], LY367385 [100 μM], picrotoxin [50 μM], CGP5462 [1 μM], atropine [50 μM], CPT [10 μM], suramin [100 μM], and flupenthixol [30 μM]). Prior to optogenetic stimulation, we monitored spontaneous calcium events in astrocytes. After 60 seconds of baseline activity, the optogenetic stimulation led to an increase in astrocyte calcium activity (Fig. 2C and D). Furthermore, we observed that optogenetic activation of CA1 neurons led to an increase in the probability of calcium events both in the somas and processes of astrocytes (Somas: from 106.25 ± 8.87 to 270.83 ± 52.49% of baseline, n=8, *p<0.05. Processes: from 86.97 ± 11.04 to 303.54 ± 34.93% of baseline, n=8, **p<0.01; Fig. 2E). Importantly, this increase was abolished when the inverse agonist of CB1R, AM-251, was present (from 82.64 ± 10.78 to 98.96 ± 21.27% of baseline, n=5, p=0.52; Fig. 2F; AM-251). Additionally, in mice lacking astroglial CB1R (GFAP-CB1R-/-), the calcium elevations in response to optogenetic stimulation were absent, while in GFAP-CB1R+/+ mice (GFAP-CB1R+/+; wild type), the calcium elevations were still observed (from 107.90 ± 6.67 to 272.01 ± 51.21% of baseline, n=9, ***p<0.001; Fig. 2F; GFAP-CB1R+/+). This suggests that astrocyte CB1Rs are responsible for mediating the astrocyte calcium mobilization in response to optogenetic stimulation of CA1 pyramidal neurons. To further investigate the mechanism of eCB-mediated calcium mobilization, we conducted optostimulation experiments using pharmacological and transgenic mouse approaches. In the presence of the synthesis enzyme antagonist of 2-AG, THL, an increase in calcium elevations was observed (from 99.68 ± 13.10 to 252.01 ± 29.27% of baseline, n=6, *p<0.05; Fig. 2F; THL). Similarly, in DAGLα-/-mice (mice lacking the DAGLα enzyme responsible for 2-AG synthesis), calcium elevations were observed (from 88.33 ± 17.92 to 274.20 ± 38.69% of baseline, n=6, **p<0.01 for DAGLα-/-; Fig. 2F; DAGLα-/-). However, when we used the antagonist LEI-401 (which inhibits the synthesis enzyme of AEA) or employed NAPE-PLD-/-mice (mice lacking the NAPE-PLD enzyme responsible for AEA synthesis), the optogenetic stimulation-induced calcium elevations were abolished (LEI-401: from 89.64 ± 8.59 to 86.23 ± 20.68% of baseline, n=10, p=0.99; Fig. 2F; LEI-401; NAPE-PLD-/-: from 106.3 ± 24.21 to 84.38 ± 32.02% of baseline, n=6, p=0.99 for NAPE-PLD-/-; Fig. 2F; NAPE-PLD-/-). This indicates that the synthesis of AEA, induced by neuronal optogenetic stimulation, is responsible for the mobilization of intracellular calcium in astrocytes. Consequently, the activation of neurons induces calcium elevations in astrocytes. However, these elevations are inhibited when the astrocyte CB1R is non-functional or when the synthesis enzyme for AEA, NAPE-PLD, is absent.

**Figure 2.**
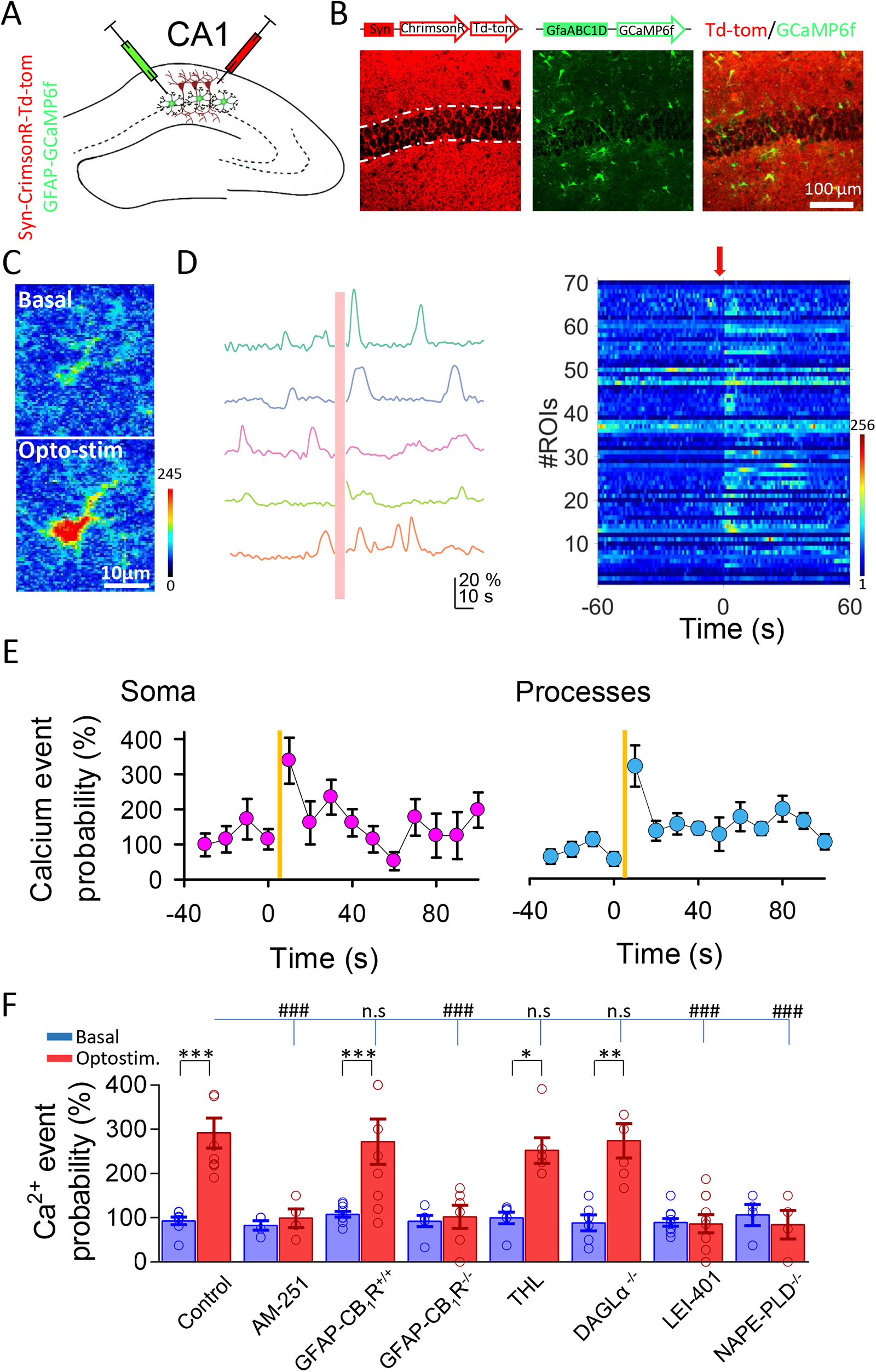
AEA released by optogenetic stimulation induces astrocyte Ca2+ mobilization. **(a)** Schematic representation of viral vectors injected into the CA1 of the hippocampus. **(b)** Fluorescence image showing ChrimsonR expression in neurons (left image), GCaMP6f in astrocytes (middle image) and merge (right image) in the CA1 of the hippocampus. (**c**) Pseudocolor images showing fluorescence intensities in CA1 astrocytes before and after ND. Scale bar, 10 µm. (**d**) left, representative traces of astrocytic calcium levels before and after optogenetic stimulation (red bar). Scale bars, 20% and 10 s. Right, heat map showing Ca^2+^-based fluorescence levels showing Ca^2+^ events to optogenetic stimulation (red arrow), in control conditions. (**e**) Relative calcium event probability before and after optogenetic stimulation at time 0 in somas and processes in control conditions. (**f**) Relative calcium event probability before and after optogenetic stimulation in control conditions, AM251, GFAP-CB1R^+/+^, GFAP-CB1R^-/-^, THL, DAGLα^-/-^, LEI401 and NAPE-PLD^-/-^.

### AEA Mediates Long-Term Synaptic Potentiation (LTP) Induced by STDP

Spike timing-dependent plasticity (STDP) is a biological process that occurs when there are precise temporal correlations between the spikes of pre- and postsynaptic neurons. This process adjusts the strength of connections between neurons in the brain and is believed to play a crucial role in learning and information storage (Feldman, 2012). Traditionally, it has been understood that NMDARs (N-methyl-D-aspartate receptors) in the postsynaptic neurons act as coincidence detectors, and their involvement in STDP has been extensively studied from both the pre- and postsynaptic perspectives (Feldman, 2012) (Supplementary Fig. 3; D-AP5). However, more recent research has demonstrated that metabotropic glutamate receptors (mGluRs) also function as detectors of coincident activity, particularly during repetitive pairings that occur within a specific time window (5-10 ms), similar to the time frame of STDP (Feldman, 2012; Markram et al., 2012) (Supplementary Fig. 3; MPEP+LY).

To investigate the involvement of 2-AG and AEA in synaptic plasticity mediated by spike timing-dependent plasticity (STDP), we conducted experiments using an STDP protocol. This protocol consisted of a subthreshold postsynaptic potential (PSP) followed by a backpropagating action potential (AP) with a delay of 10 ms. The protocol was repeated 100 times at a frequency of 0.2 Hz in CA1 pyramidal neurons of the hippocampus (Fig. 3A and B). Under control conditions, the 100 pairings of the STDP protocol induced long-term potentiation (LTP) (from 101.20 ± 2.31 to 161.40 ± 15.18% of peak amplitude, n=8, ***p<0.001; Fig. 3C, D, and G; control).

**Figure 3.**
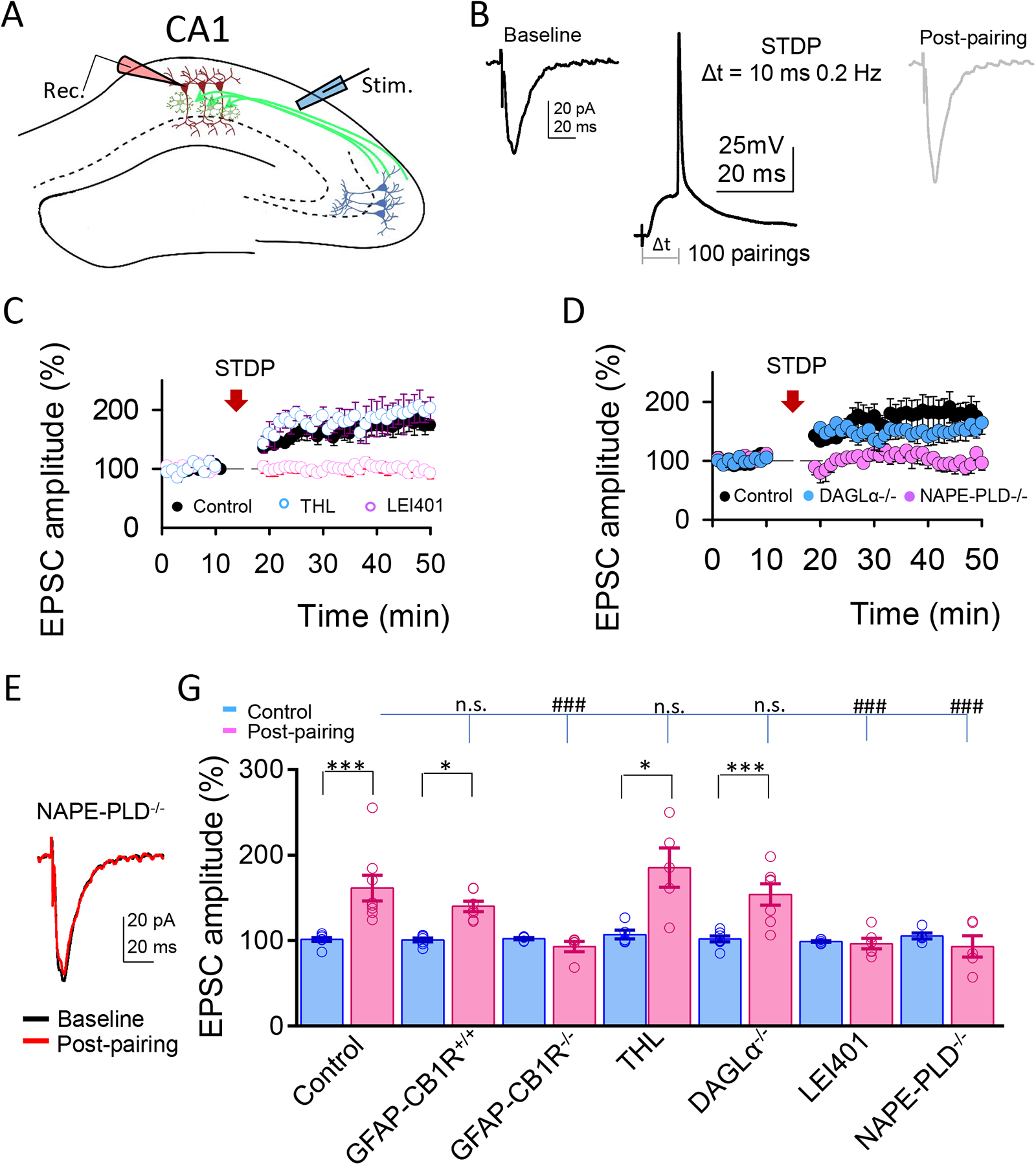
AEA-Induced astrocyte activation is necessary for LTP induced by STDP. (**a**) Schematic representation of the experimental design. (**b**) *left*. Representative EPSCs before (left) and after (right) PSP followed by an AP with a 10 ms delay and a frequency of 0.2 Hz (middle). (**c**) Time course of EPSC amplitude before and after 100 pairing for STDP in control (black circles), in the presence of THL (empty blue circles) and in the presence of LEI401 (empty pink circles). (**d**) Time course of EPSC amplitude before and after 100 pairing for STDP in control (black circles), in DAGLα^-/-^ (blue circles) and in NAPE-PLD^-/-^ (pink circles). (**e**) *left*. Representative EPSCs traces before (black trace) and after (red trace) 100 pairing for STDP in NAPE-PLD^-/-^ mice. (**f**) EPSC amplitude during baseline (blue bars) and after 100 pairings (pink bars) in different experimental conditions: Control, IP3R2^-/-,^ GFAP-CB1R^+/+^, GFAP-CB1R^-/-,^ THL, DAGLα^-/-^, LEI401, NAPE-PLD^-/-^.

Previous studies have demonstrated that eCB signaling to astrocytes plays a crucial role in controlling synaptic plasticity induced by spike timing-dependent plasticity (STDP) in different brain areas (Andrade-Talavera et al., 2016; Falcón-Moya et al., 2020; Min & Nevian, 2012). To investigate the involvement of astrocytes in STDP-induced long-term potentiation (LTP), we employed various experimental approaches. First, we utilized the calcium chelator BAPTA in the internal solution of a single astrocyte. BAPTA can diffuse through the extensive network of astrocytes connected by gap junctions (Covelo & Araque, 2018; Navarrete & Araque, 2010a; Perea & Araque, 2007). After 20-30 minutes of BAPTA diffusion, the induction of LTP failed to occur (from 94.68 ± 1.64 to 93.88 ± 15.27% of peak amplitude, n=6, p=0.99; Supplementary Fig. 4; aBAPTA). Additionally, LTP was absent in mice lacking IP3 receptor type 2 (from 100.90 ± 2.05 to 109.70 ± 8.69% of peak amplitude, n=6, p=0.99; Supplementary Fig. 4; IP3R2-/-), indicating the essential role of astrocytes in LTP induction. Next, we investigated the involvement of astrocyte CB1R by conducting experiments in mice lacking astroglial CB1R. While LTP was still observed in GFAP-CB1R+/+ mice (from 101.30 ± 1.58 to 150.8 ± 9.58% of peak amplitude, n=7, *p<0.05; Fig. 3G, GFAP-CB1R+/+), it was abolished in GFAP-CB1R-/-mice (from 102.20 ± 1.17 to 92.98 ± 6.12% of peak amplitude, n=5, p=0.99; Fig. 3G, GFAP-CB1R-/-), demonstrating the essential role of astroglial CB1R in LTP induction. Furthermore, we investigated the specific eCB involved in LTP by blocking the synthesis enzyme for 2-AG, DAGLα, using the THL antagonist. The inhibition of DAGLα with THL did not affect the induction of LTP (from 107.01 ± 5.31 to 185.20 ± 22.96% of peak amplitude, n=5, ***p<0.001; Fig. 3C and G, THL). Similarly, LTP was still observed in mice lacking DAGLα (from 101.70 ± 3.47 to 153.90 ± 12.51% of peak amplitude, n=7, *p<0.05; Fig. 3D and G, DAGLα-/-), indicating that 2-AG synthesis is not necessary for LTP induction. The blockade of the enzyme responsible for synthesizing AEA, NAPE-PLD, using LEI-401 or in NAPE-PLD knockout mice, resulted in the prevention of LTP induction (LEI-401: from 98.60 ± 0.95 to 96.44 ± 6.02% of peak amplitude, n=6, p=0.99; Fig. 3C and G, LEI401; NAPE-PLD-/-: from 105.30 ± 3.44 to 93.04 ± 12.62% of peak amplitude, n=5, p=0.99; Fig. 3D, E and G, NAPE-PLD-/-). These findings suggest that AEA signaling is crucial for this form of neuronal plasticity. Taken together, the results demonstrate that AEA synthesis through astrocyte CB1R activation plays a critical role in the induction of LTP in CA1 pyramidal neurons of the hippocampus. These findings highlight the importance of eCB signaling and astrocyte involvement in synaptic plasticity processes such as STDP-induced LTP.

## Discussion

Using a combination of two-photon imaging and pair-recordings of CA1 pyramidal neurons, employing a minimal stimulation technique, our research has uncovered an unexpected finding. We have observed that the release of endocannabinoids (eCB) induces endocannabinoid-mediated synaptic potentiation (e-SP) through the involvement of astrocytes, even in synapses located at a considerable distance from the source (Covelo & Araque, 2018; Martín et al., 2015; Martin-Fernandez et al., 2017a; Navarrete & Araque, 2010a). Our investigations revealed that the occurrence of e-SP was effectively prevented when the NAPE-PLD enzyme, responsible for eCB synthesis, was pharmacologically or genetically deleted, as well as in the absence of astroglial CB1 receptors. These results strongly indicate that anandamide (AEA) serves as the specific eCB responsible for signaling to astrocytes. Interestingly, the e-SP phenomenon persisted despite the pharmacological or genetic blockade of the synthesis enzyme for 2-arachidonoylglycerol (2-AG), DAGLα. Moreover, we observed that depolarization-induced suppression of excitation (DSE) was abolished when DAGLα activity was either pharmacologically or genetically prevented, suggesting that 2-AG specifically signals to neurons, inducing DSE. Furthermore, our findings demonstrated that only the inactivation of NAPE-PLD or the deletion of astroglial CB1 receptors prevented the mobilization of intracellular calcium in astrocytes in response to eCB release induced by neuronal activation. This implies a distinctive signaling pattern for 2-AG and AEA, resulting in contrasting synaptic regulation mediated by the same neuronal and astroglial CB1 receptors.

Lastly, we discovered that astrocyte-driven synaptic plasticity relies on AEA synthesis and functional astrocytic CB1 receptors for long-term potentiation (LTP) induced by spike-timing-dependent plasticity (STDP). These results provide compelling evidence that 2-AG and AEA induce distinct and contrasting synaptic regulations through the same CB1 receptors across different cell types.

The findings presented in this study challenge the conventional understanding of neurotransmitter systems, which typically involve one principal agonist acting on multiple receptor types. In contrast, our research demonstrates that the endocannabinoid (eCB) system employs a distinctive signaling mechanism, utilizing the same CB1 receptor but with different agonists, AEA and 2-AG, to selectively communicate with either astrocytes or neurons. This expands our understanding of neuromodulation in the brain.

Previous studies have established that eCBs exert their inhibitory effects over short distances (<20 μm), as documented in research by Chevaleyre et al. (2006), Chevaleyre & Castillo (2004), Piomelli (2003), Wilson & Nicoll (2002). Additionally, eCBs can also transmit signals over long distances through the spreading of astrocytic calcium signals, subsequently leading to the release of gliotransmitters in more distant synapses (Covelo & Araque, 2018; Martín et al., 2015; Martin-Fernandez et al., 2017a; Navarrete & Araque, 2010a). However, the specific eCB responsible for both phenomena has not been thoroughly investigated. In this study, we provide the description of 2-AG signaling to neurons, inducing depolarization-induced suppression of excitation (DSE) through presynaptic CB1 receptors. Simultaneously, AEA selectively signals to astrocytes, triggering endocannabinoid-mediated synaptic potentiation (e-SP) via astroglial CB1 receptors and subsequent mobilization of astrocytic calcium. Overall, our findings challenge the traditional notion of neurotransmitter systems and demonstrate the intricate and distinct roles played by AEA and 2-AG in modulating neuronal and astrocytic signaling through the CB1 receptor.

The involvement of 2-AG in both short-term and long-term synaptic plasticity, regulated exclusively through neuronal mechanisms, has been extensively studied and well-documented across various brain regions (Castillo et al., 2012; Kano et al., 2009; Katona & Freund, 2012; Ohno-Shosaku et al., 2012; Piomelli, 2014). Previous research has demonstrated that depolarization-induced suppression of inhibition (DSI), depolarization-induced suppression of excitation (DSE), and endocannabinoid-induced short-term potentiation (STP) are all inhibited by pharmacological and genetic deletion of DAGLα, the enzyme responsible for 2-AG synthesis (Gao et al., 2010; Hashimotodani et al., 2013; Tanimura et al., 2010; Yoshino et al., 2011). Conversely, studies have shown that blocking FAAH, the degradation enzyme of AEA, does not impact these synaptic plasticity mechanisms (Hashimotodani et al., 2007). These previous findings align with our present data, which indicate that the deletion of DAGLα, the synthesis enzyme of 2-AG, abolishes DSE. Furthermore, we demonstrate that blocking NAPE-PLD or using mice lacking astroglial CB1 receptors in the hippocampus does not affect synaptic transmission.

While 2-AG is widely recognized as the primary player in most forms of homo- and heterosynaptic short-term depression, as well as certain forms of long-term depression (Heifets & Castillo, 2009; Kano et al., 2009), AEA has also been shown to act through CB1 receptors in various neurobiological contexts (Clapper et al., 2010; Kinsey et al., 2009; Straiker et al., 2011). AEA mediates specific forms of synaptic regulation through presynaptic CB1 receptors (Ade & Lovinger, 2007; Adermark & Lovinger, 2007; Gerdeman et al., 2002; Kim & Alger, 2010). Our research, employing pair-recordings of CA1 pyramidal neurons and a minimal stimulation technique, has uncovered that the release of eCBs by neurons induces endocannabinoid-mediated synaptic potentiation (e-SP) through the involvement of astrocytes, even in distant synapses (Covelo & Araque, 2018; Martín et al., 2015; Martin-Fernandez et al., 2017a; Navarrete & Araque, 2010a). We have established that the occurrence of e-SP is prevented when NAPE-PLD, the enzyme responsible for AEA synthesis, is pharmacologically or genetically deleted, or when astroglial CB1 receptors are absent. This indicates that AEA specifically signals to astrocytes, inducing e-SP. However, the e-SP phenomenon persists despite pharmacological and genetic blockade of DAGLα, the enzyme involved in 2-AG synthesis. This suggests a differential signaling mechanism for 2-AG and AEA, resulting in contrasting synaptic regulation mediated by the same neuronal and astroglial CB1 receptors.

The involvement of endocannabinoid (eCB) signaling to astrocytes in certain forms of synaptic plasticity, such as spike-timing dependent plasticity (STDP), holds significant implications. Traditionally, eCBs were primarily studied for their canonical retrograde signaling role in neuronal communication (Markram et al., 2011). However, the emerging understanding of eCB signaling to astrocytes in this specific type of synaptic regulation highlights the crucial involvement of astrocyte activation in the induction of synaptic plasticity underlying associative learning and information storage in the brain (Andrade-Talavera et al., 2016; Falcón-Moya et al., 2020; Min & Nevian, 2012). Our research findings demonstrate the critical role of anandamide (AEA) in the induction of long-term potentiation (LTP) triggered by STDP. Furthermore, the functionality of astroglial CB1 receptors was found to be decisive for this particular form of synaptic plasticity. The distinct roles of AEA and 2-AG may be reflected in their functional or spatial segregation within the brain. Thus, by acting from different synapses and influencing distinct microcircuits, AEA and 2-AG can be considered specialized neuromodulators, redefining our understanding of neuromodulation.

To summarize, our findings demonstrate that 2-AG predominantly signals to neurons and leads to a decrease in synaptic efficacy through exclusively neuronal mechanisms. On the other hand, AEA primarily signals to astrocytes and enhances synaptic efficacy through the activation of astrocyte CB1 receptors. The cumulative results emphasize that different endocannabinoids exhibit distinct signaling patterns to astrocytes and neurons, resulting in diverse and contrasting forms of synaptic regulation. These findings challenge the traditional perspective of neurotransmitter systems and pave the way for a new conceptualization of neuromodulation in brain function.

## Figure legends

**Supplementary Figure 1.**
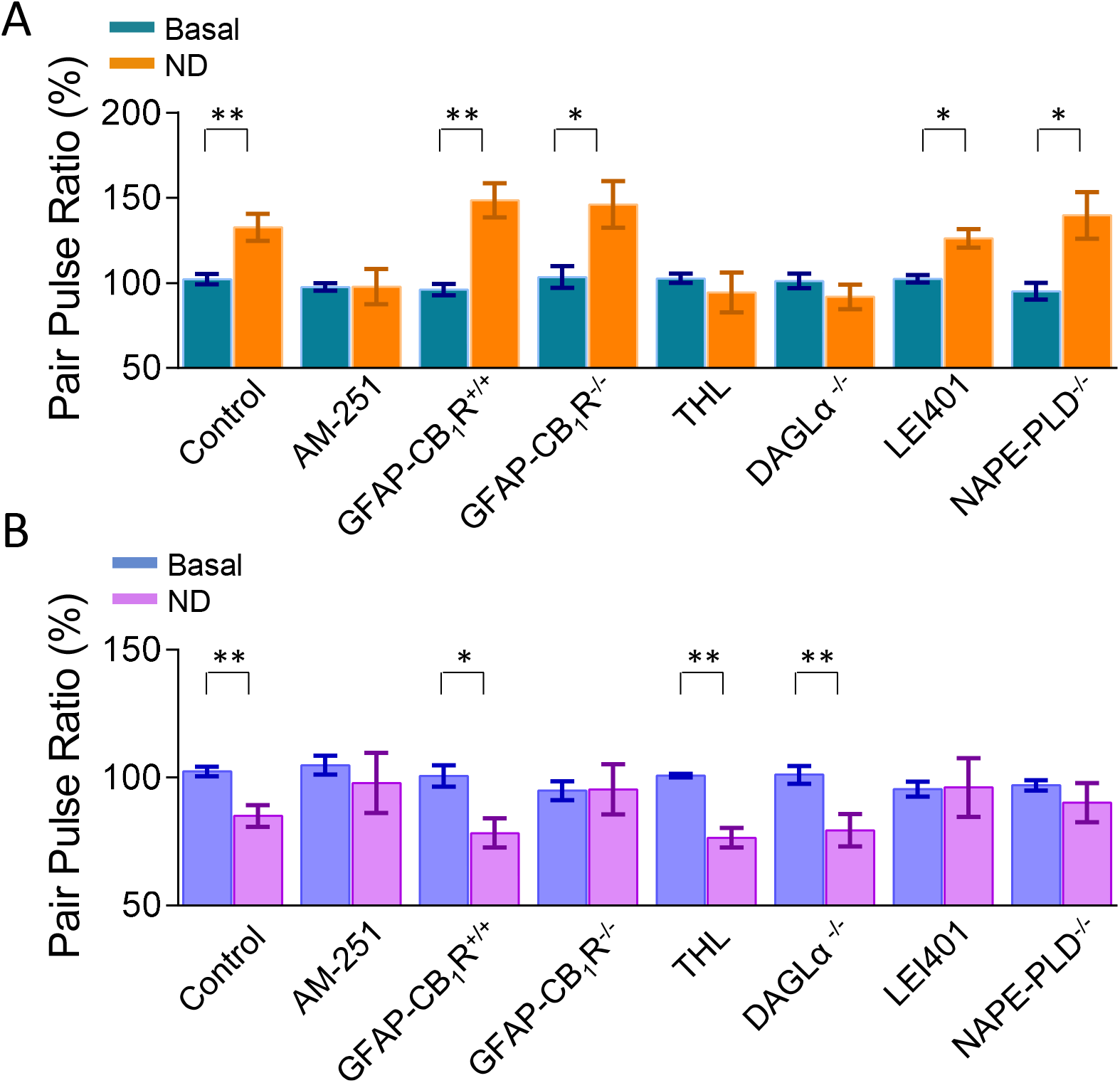
Pair pulse ratio of EPSC from CA1 pyramidal neurons in different experimental conditions. (a) Pair pulse ratio of the EPSCs before and after homoneuron ND in control conditions, AM251, GFAP-CB1R^+/+^, GFAP-CB1R^-/-^, THL, DAGLα^-/-^, LEI401, NAPE-PLD^-/-^. (**a**) Pair pulse ratio of the EPSCs before and after heteroneuron ND in control conditions, AM251, GFAP-CB1R^+/+^, GFAP-CB1R^-/-^, THL, DAGLα^-/-^, LEI401, NAPE-PLD^-/-^.

**Supplementary Figure 2.**
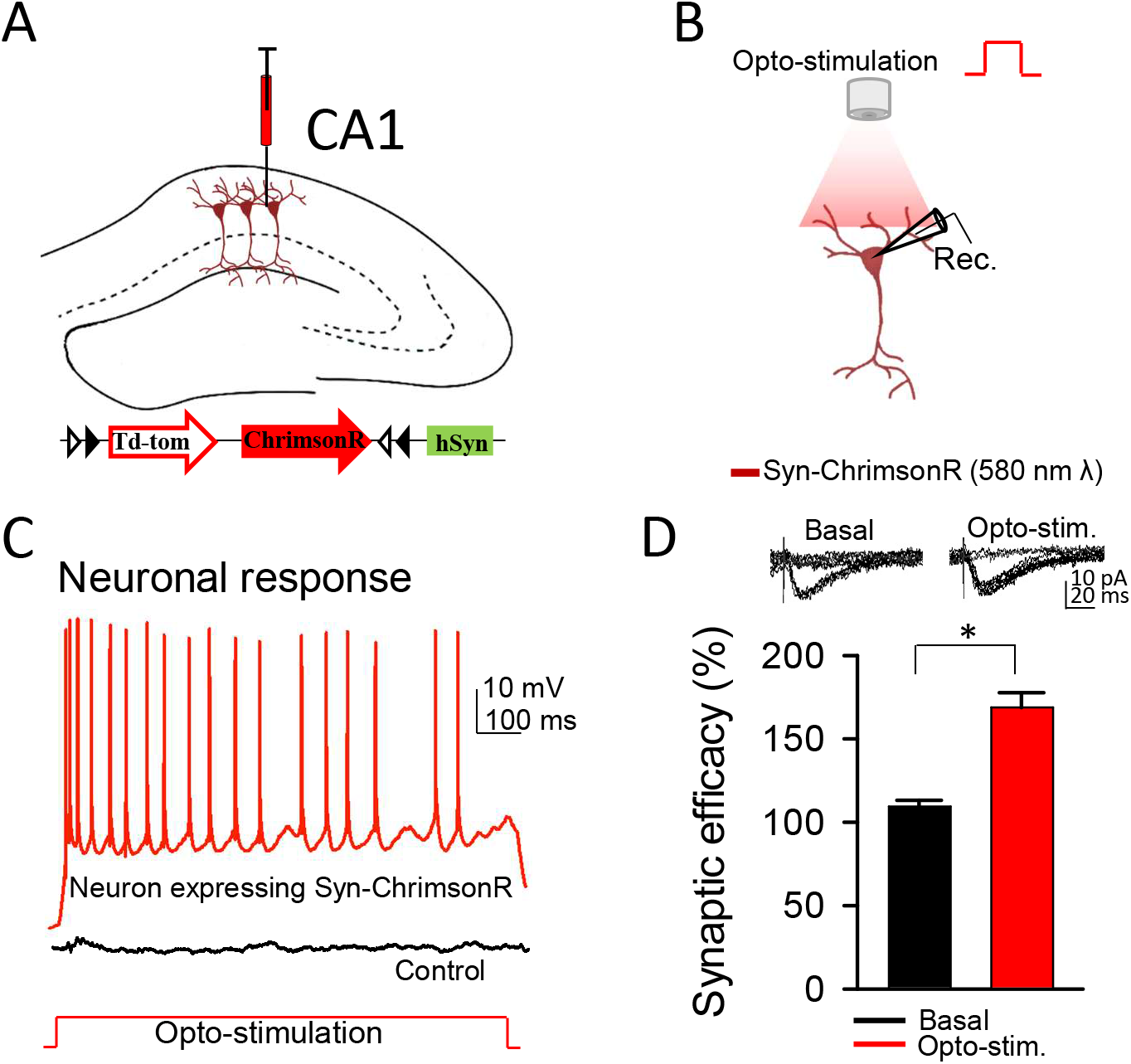
Enhanced synaptic efficacy by optogenetic stimulation of CA1 pyramidal neurons. (**a**) Drawing showing viral vectors injected into the CA1 of the hippocampus. (**b**) schematic representation of the experimental design. (**c**) Neuronal response to optogenetic stimulation of CA1 pyramidal neuron expressing the opsin crimsonR (red trace) and empty virus (black trace). (**d**) Synaptic efficacy of EPSC during baseline (black bar) and after opto-stimulation (red bar).

**Supplementary Figure 3.**
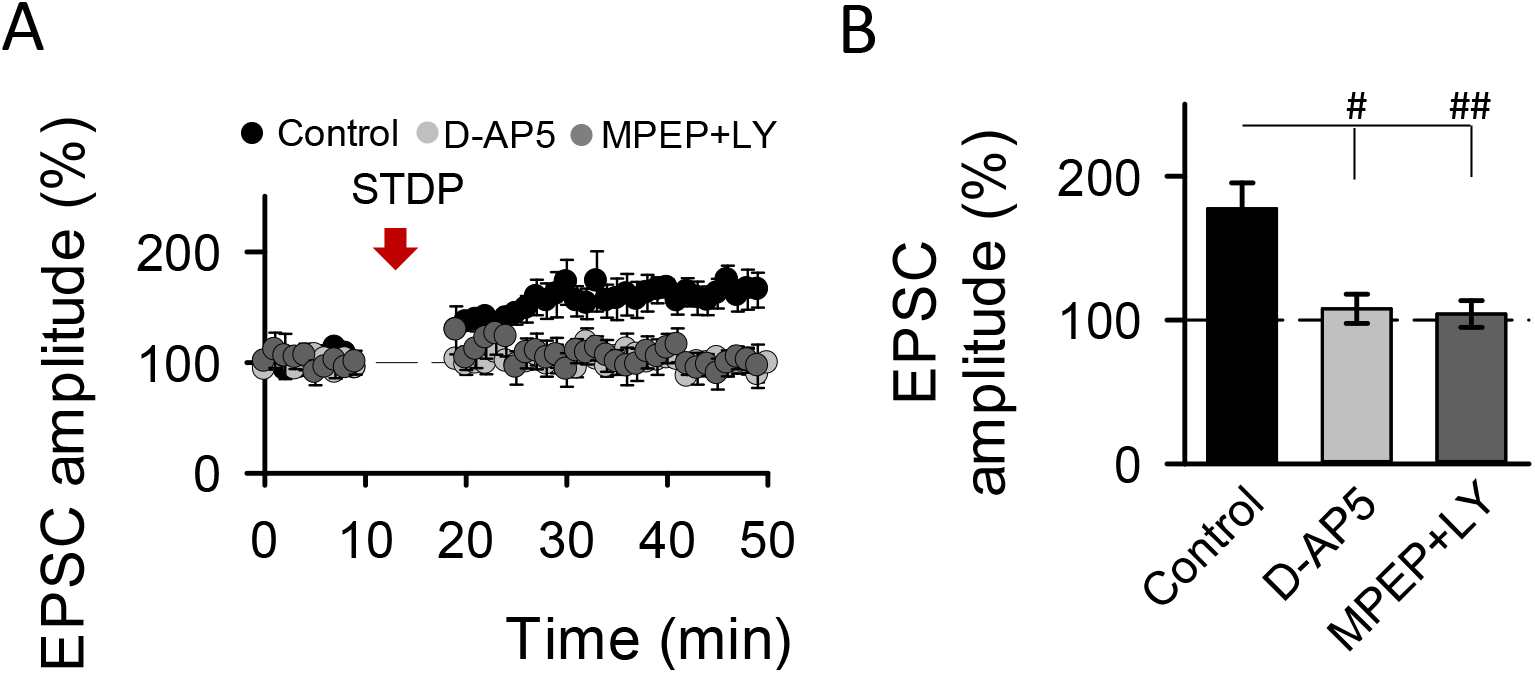
STDP-induced LTP depends on NMDA and Group I mGluR. **(a)** Time course of the LTP induced by STDP in control condition (black circles; control), under NMDAR antagonist, D-AP5 (light grey circles; D-AP5), and in the presence of group I mGluR antagonists, MPEP and LY (dark grey circles; MPEP + LY). **(b)** Bargraph of the LTP induced by STDP in control condition (black graph; control), under NMDAR antagonist, D-AP5 (light grey graph; D-AP5) and in the presence of group I mGluR antagonists, MPEP and LY (dark grey graph; MPEP + LY). Note that LTP is abolished in the presence of D-AP5 and MPEP + LY, demonstrating that LTP depends on NMDAR and Group I mGluRs.

**Supplementary Figure 4.**
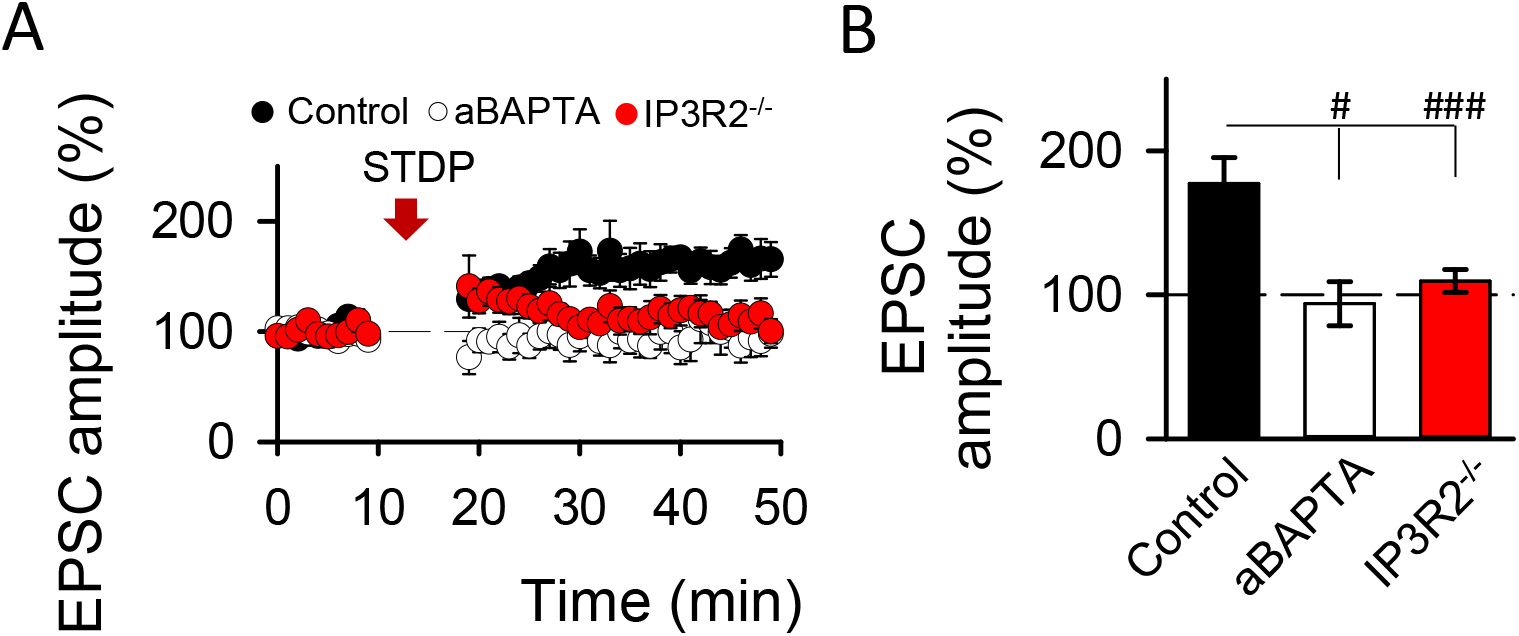
STDP-induced LTP depends on intracellular calcium mobilization in astrocytes. **(a)** Time course of the LTP induced by STDP in control condition (black circles; control), in calcium chelator BAPTA loading astrocytes (white circles; aBAPTA), and in IP3R2-/-(red circles; IP3R2-/-). **(b)** Bargraph of the LTP induced by STDP in control condition (black circles; control), in calcium chelator BAPTA loading astrocytes (white circles; aBAPTA), and in IP3R2-/-(red circles; IP3R2-/-). Note that LTP is prevented in BAPTA loading astrocyte network and in IP3R2-/-, demonstrating the necessary role of astrocytes activation LTP induction.

## Bibliography

Ade, K. K., & Lovinger, D. M. (2007). Anandamide regulates postnatal development of long-term synaptic plasticity in the rat dorsolateral striatum. The Journal of Neurosciencel: The Official Journal of the Society for Neuroscience, 27(9), 2403–2409. https://doi.org/10.1523/JNEUROSCI.2916-06.2007

Adermark, L., & Lovinger, D. M. (2007). Retrograde endocannabinoid signaling at striatal synapses requires a regulated postsynaptic release step. Proceedings of the National Academy of Sciences of the United States of America, 104(51), 20564–20569. https://doi.org/10.1073/PNAS.0706873104

Ahn, K., McKinney, M. K., & Cravatt, B. F. (2008). Enzymatic Pathways That Regulate Endocannabinoid Signaling in the Nervous System. Chemical Reviews, 108(5), 1687– 1707. https://doi.org/10.1021/cr0782067

Alger, B. E., & Kim, J. (2011). Supply and demand for endocannabinoids. Trends in Neurosciences, 34(6), 304–315. https://doi.org/10.1016/J.TINS.2011.03.003

Andrade-Talavera, Y., Duque-Feria, P., Paulsen, O., & Rodríguez-Moreno, A. (2016). Presynaptic Spike Timing-Dependent Long-Term Depression in the Mouse Hippocampus. Cerebral Cortex, 26(8), 3637–3654. https://doi.org/10.1093/cercor/bhw172

Baraibar, A. M., Belisle, L., Marsicano, G., Matute, C., Mato, S., Araque, A., & Kofuji, P. (2022). Spatial Organization of Neuron-Astrocyte Interactions in the Somatosensory Cortex. BioRxiv, 2022.02.03.479046. https://doi.org/10.1101/2022.02.03.479046

Castillo, P. E., Younts, T. J., Chávez, A. E., & Hashimotodani, Y. (2012). Endocannabinoid Signaling and Synaptic Function. In Neuron. https://doi.org/10.1016/j.neuron.2012.09.020

Chevaleyre, V., & Castillo, P. E. (2004). Endocannabinoid-mediated metaplasticity in the hippocampus. Neuron, 43(6), 871–881. https://doi.org/10.1016/j.neuron.2004.08.036

Chevaleyre, V., Takahashi, K. A., & Castillo, P. E. (2006). Endocannabinoid-Mediated Synaptic Plasticity in the CNS. https://doi.org/10.1146/annurev.neuro.29.051605.112834

Clapper, J. R., Moreno-Sanz, G., Russo, R., Guijarro, A., Vacondio, F., Duranti, A., Tontini, A., Sanchini, S., Sciolino, N. R., Spradley, J. M., Hohmann, A. G., Calignano, A., Mor, M., Tarzia, G., & Piomelli, D. (2010). Anandamide suppresses pain initiation through a peripheral endocannabinoid mechanism. Nature Neuroscience, 13(10), 1265– 1270. https://doi.org/10.1038/NN.2632

Covelo, A., & Araque, A. (2018). Neuronal activity determines distinct gliotransmitter release from a single astrocyte. ELife, 7, e32237. https://doi.org/10.7554/eLife.32237

Covelo, A., Eraso-Pichot, A., Fernández-Moncada, I., Serrat, R., & Marsicano, G. (2021). CB1R-dependent regulation of astrocyte physiology and astrocyte-neuron interactions. Neuropharmacology, 195, 108678. https://doi.org/10.1016/J.NEUROPHARM.2021.108678

Cravatt, B. F., Giang, D. K., Mayfield, S. P., Boger, D. L., Lerner, R. A., & Gilula, N. B. (1996). Molecular characterization of an enzyme that degrades neuromodulatory fatty-acid amides. Nature, 384(6604), 83–87. https://doi.org/10.1038/384083A0

Di Marzo, V. (2009). The endocannabinoid system: Its general strategy of action, tools for its pharmacological manipulation and potential therapeutic exploitation. In Pharmacological Research. https://doi.org/10.1016/j.phrs.2009.02.010

Dinh, T. P., Freund, T. F., & Piomelli, D. (2002). A role for monoglyceride lipase in 2-arachidonoylglycerol inactivation. Chemistry and Physics of Lipids, 121(1–2), 149–158. https://doi.org/10.1016/S0009-3084(02)00150-0

Dobrunz, L. E., & Stevens, C. F. (1997). Heterogeneity of release probability, facilitation, and depletion at central synapses. Neuron, 18(6), 995–1008. https://doi.org/10.1016/S0896-6273(00)80338-4

Falcón-Moya, R., Pérez-Rodríguez, M., Prius-Mengual, J., Andrade-Talavera, Y., Arroyo-García, L. E., Pérez-Artés, R., Mateos-Aparicio, P., Guerra-Gomes, S., Oliveira, J. F., Flores, G., & Rodríguez-Moreno, A. (2020). Astrocyte-mediated switch in spike timing-dependent plasticity during hippocampal development. Nature Communications. https://doi.org/10.1038/s41467-020-18024-4

Feldman, D. E. (2012). The spike timing dependence of plasticity. Neuron, 75(4), 556. https://doi.org/10.1016/J.NEURON.2012.08.001

Gao, Y., Vasilyev, D. V., Goncalves, M. B., Howell, F. V., Hobbs, C., Reisenberg, M., Shen, R., Zhang, M. Y., Strassle, B. W., Lu, P., Mark, L., Piesla, M. J., Deng, K., Kouranova, E. V., Ring, R. H., Whiteside, G. T., Bates, B., Walsh, F. S., Williams, G., … Doherty, P. (2010). Loss of retrograde endocannabinoid signaling and reduced adult neurogenesis in diacylglycerol lipase knock-out mice. The Journal of Neurosciencel: The Official Journal of the Society for Neuroscience, 30(6), 2017–2024. https://doi.org/10.1523/JNEUROSCI.5693-09.2010

Gerdeman, G. L., Ronesi, J., & Lovinger, D. M. (2002). Postsynaptic endocannabinoid release is critical to long-term depression in the striatum. Nature Neuroscience, 5(5), 446–451. https://doi.org/10.1038/NN832

Gómez-Gonzalo, M., Navarrete, M., Perea, G., Covelo, A., Martín-Fernández, M., Shigemoto, R., Luján, R., & Araque, A. (2015). Endocannabinoids induce lateral long-term potentiation of transmitter release by stimulation of gliotransmission. Cerebral Cortex, 25(10), 3699–3712. https://doi.org/10.1093/cercor/bhu231

Hashimotodani, Y., Ohno-Shosaku, T., & Kano, M. (2007). Presynaptic monoacylglycerol lipase activity determines basal endocannabinoid tone and terminates retrograde endocannabinoid signaling in the hippocampus. The Journal of Neurosciencel: The Official Journal of the Society for Neuroscience, 27(5), 1211–1219. https://doi.org/10.1523/JNEUROSCI.4159-06.2007

Hashimotodani, Y., Ohno-Shosaku, T., Tanimura, A., Kita, Y., Sano, Y., Shimizu, T., Di Marzo, V., & Kano, M. (2013). Acute inhibition of diacylglycerol lipase blocks endocannabinoid-mediated retrograde signalling: evidence for on-demand biosynthesis of 2-arachidonoylglycerol. The Journal of Physiology, 591(19), 4765–4776. https://doi.org/10.1113/JPHYSIOL.2013.254474

Heifets, B. D., & Castillo, P. E. (2009). Endocannabinoid Signaling and Long-Term Synaptic Plasticity. Annual Review of Physiology, 71(1), 283–306. https://doi.org/10.1146/annurev.physiol.010908.163149

Hillard, C. J., Wilkison, D. M., Edgemond, W. S., & Campbell, W. B. (1995). Characterization of the kinetics and distribution of N-arachidonylethanolamine (anandamide) hydrolysis by rat brain. Biochimica et Biophysica Acta, 1257(3), 249–256. https://doi.org/10.1016/0005-2760(95)00087-S

Isaac, J. T. R., Hjelmstad, G. O., Nicoll, R. A., & Malenka, R. C. (1996). Long-term potentiation at single fiber inputs to hippocampal CA1 pyramidal cells. Proceedings of the National Academy of Sciences, 93(16), 8710–8715. https://doi.org/10.1073/PNAS.93.16.8710

Kano, M., Ohno-Shosaku, T., Hashimotodani, Y., Uchigashima, M., & Watanabe, M. (2009). Endocannabinoid-mediated control of synaptic transmission. In Physiological Reviews. https://doi.org/10.1152/physrev.00019.2008

Katona, I., & Freund, T. F. (2012). Multiple functions of endocannabinoid signaling in the brain. Annual Review of Neuroscience, 35, 529–558. https://doi.org/10.1146/ANNUREV-NEURO-062111-150420

Kim, J., & Alger, B. E. (2010). Reduction in endocannabinoid tone is a homeostatic mechanism for specific inhibitory synapses. Nature Neuroscience, 13(5), 592–600. https://doi.org/10.1038/NN.2517

Kinsey, S. G., Long, J. Z., O’Neal, S. T., Abdullah, R. A., Poklis, J. L., Boger, D. L., Cravatt, B. F., & Lichtman, A. H. (2009). Blockade of endocannabinoid-degrading enzymes attenuates neuropathic pain. The Journal of Pharmacology and Experimental Therapeutics, 330(3), 902–910. https://doi.org/10.1124/JPET.109.155465

Klapoetke, N. C., Murata, Y., Kim, S. S., Pulver, S. R., Birdsey-Benson, A., Cho, Y. K., Morimoto, T. K., Chuong, A. S., Carpenter, E. J., Tian, Z., Wang, J., Xie, Y., Yan, Z., Zhang, Y., Chow, B. Y., Surek, B., Melkonian, M., Jayaraman, V., Constantine-Paton, M., … Boyden, E. S. (2014). Independent optical excitation of distinct neural populations. Nature Methods, 11(3), 338–346. https://doi.org/10.1038/nmeth.2836

Marcus, D. J., Bedse, G., Gaulden, A. D., Ryan, J. D., Kondev, V., Winters, N. D., Rosas-Vidal, L. E., Altemus, M., Mackie, K., Lee, F. S., Delpire, E., & Patel, S. (2020). Endocannabinoid Signaling Collapse Mediates Stress-Induced Amygdalo-Cortical Strengthening. Neuron, 105(6), 1062. https://doi.org/10.1016/J.NEURON.2019.12.024

Markram, H., Gerstner, W., & Sjöström, P. J. (2011). A History of Spike-Timing-Dependent Plasticity. Frontiers in Synaptic Neuroscience, 3(AUG), 1–24. https://doi.org/10.3389/FNSYN.2011.00004

Markram, H., Gerstner, W., & Sjöström, P. J. (2012). Spike-timing-dependent plasticity: a comprehensive overview. Frontiers in Synaptic Neuroscience, 4(JULY). https://doi.org/10.3389/FNSYN.2012.00002

Martín, R., Bajo-Grañeras, R., Moratalla, R., Perea, G., & Araque, A. (2015). Circuit-specific signaling in astrocyte-neuron networks in basal ganglia pathways. Science (New York, N.Y.), 349(6249), 730–734. https://doi.org/10.1126/SCIENCE.AAA7945

Martin-Fernandez, M., Jamison, S., Robin, L. M., Zhao, Z., Martin, E. D., Aguilar, J., Benneyworth, M. A., Marsicano, G., & Araque, A. (2017a). Synapse-specific astrocyte gating of amygdala-related behavior. Nature Neuroscience, 20(11), 1540–1548. https://doi.org/10.1038/nn.4649

Martin-Fernandez, M., Jamison, S., Robin, L. M., Zhao, Z., Martin, E. D., Aguilar, J., Benneyworth, M. A., Marsicano, G., & Araque, A. (2017b). Synapse-specific astrocyte gating of amygdala-related behavior. Nature Neuroscience, 20(11), 1540–1548. https://doi.org/10.1038/nn.4649

McKinney, M. K., & Cravatt, B. E. (2005). Structure and function of fatty acid amide hydrolase. Annual Review of Biochemistry, 74, 411–432. https://doi.org/10.1146/ANNUREV.BIOCHEM.74.082803.133450

Min, R., & Nevian, T. (2012). Astrocyte signaling controls spike timing-dependent depression at neocortical synapses. Nature Neuroscience, 15(5), 746–753. https://doi.org/10.1038/nn.3075

Murataeva, N., Straiker, A., & MacKie, K. (2014). Parsing the players: 2-arachidonoylglycerol synthesis and degradation in the CNS. British Journal of Pharmacology, 171(6), 1379–1391. https://doi.org/10.1111/BPH.12411

Navarrete, M., & Araque, A. (2008). Endocannabinoids Mediate Neuron-Astrocyte Communication. Neuron, 57(6), 883–893. https://doi.org/10.1016/j.neuron.2008.01.029

Navarrete, M., & Araque, A. (2010a). Endocannabinoids potentiate synaptic transmission through stimulation of astrocytes. Neuron. https://doi.org/10.1016/j.neuron.2010.08.043

Navarrete, M., & Araque, A. (2010b). Endocannabinoids potentiate synaptic transmission through stimulation of astrocytes. Neuron, 68(1), 113–126. https://doi.org/10.1016/j.neuron.2010.08.043

Noriega-Prieto, J. A., Kofuji, P., & Araque, A. (2023). Endocannabinoid signaling in synaptic function. Glia, 71(1), 36–43. https://doi.org/10.1002/GLIA.24256

Ohno-Shosaku, T., & Kano, M. (2014). Endocannabinoid-mediated retrograde modulation of synaptic transmission. Current Opinion in Neurobiology, 29, 1–8. https://doi.org/10.1016/J.CONB.2014.03.017

Ohno-Shosaku, T., Tanimura, A., Hashimotodani, Y., & Kano, M. (2012). Endocannabinoids and retrograde modulation of synaptic transmission. *The Neuroscientist*l*: A Review Journal Bringing Neurobiology*, Neurology and Psychiatry, 18(2), 119–132. https://doi.org/10.1177/1073858410397377

Perea, G., & Araque, A. (2007). Astrocytes potentiate transmitter release at single hippocampal synapses. Science, 317(5841), 1083–1086. https://doi.org/10.1126/science.1144640

Piomelli, D. (2003). The molecular logic of endocannabinoid signalling. Nature Reviews Neuroscience. https://doi.org/10.1038/nrn1247

Piomelli, D. (2014). More surprises lying ahead. The endocannabinoids keep us guessing. Neuropharmacology, 76 *Pt B*(PART B), 228–234. https://doi.org/10.1016/J.NEUROPHARM.2013.07.026

Raastad, M. (1995). Extracellular activation of unitary excitatory synapses between hippocampal CA3 and CA1 pyramidal cells. The European Journal of Neuroscience, 7(9), 1882–1888. https://doi.org/10.1111/J.1460-9568.1995.TB00709.X

Robin, L. M., Oliveira da Cruz, J. F., Langlais, V. C., Martin-Fernandez, M., Metna-Laurent, M., Busquets-Garcia, A., Bellocchio, L., Soria-Gomez, E., Papouin, T., Varilh, M., Sherwood, M. W., Belluomo, I., Balcells, G., Matias, I., Bosier, B., Drago, F., Van Eeckhaut, A., Smolders, I., Georges, F., … Marsicano, G. (2018). Astroglial CB1 Receptors Determine Synaptic D-Serine Availability to Enable Recognition Memory. Neuron. https://doi.org/10.1016/j.neuron.2018.04.034

Serrat, R., Covelo, A., Kouskoff, V., Delcasso, S., Ruiz-Calvo, A., Chenouard, N., Stella, C., Blancard, C., Salin, B., Julio-Kalajzić, F., Cannich, A., Massa, F., Varilh, M., Deforges, S., Robin, L. M., De Stefani, D., Busquets-Garcia, A., Gambino, F., Beyeler, A., … Marsicano, G. (2022). Astroglial ER-mitochondria calcium transfer mediates endocannabinoid-dependent synaptic integration. Cell Reports, 41(2), 111499. https://doi.org/10.1016/J.CELREP.2022.111499

Skupio, U., Welte, J., Serrat, R., Eraso-Pichot, A., Julio-Kalajzić, F., Gisquet, D., Cannich, A., Delcasso, S., Matias, I., Fundazuri, U. B., Pouvreau, S., Pagano Zottola, A. C., Lavanco, G., Drago, F., Ruiz de Azua, I., Lutz, B., Bellocchio, L., Busquets-Garcia, A., Chaouloff, F., & Marsicano, G. (2023). Mitochondrial cannabinoid receptors gate corticosterone impact on novel object recognition. Neuron. https://doi.org/10.1016/J.NEURON.2023.04.001

Straiker, A., Wager-Miller, J., Hu, S. S., Blankman, J. L., Cravatt, B. F., & MacKie, K. (2011). COX-2 and fatty acid amide hydrolase can regulate the time course of depolarization-induced suppression of excitation. British Journal of Pharmacology, 164(6), 1672–1683. https://doi.org/10.1111/J.1476-5381.2011.01486.X

Tanimura, A., Yamazaki, M., Hashimotodani, Y., Uchigashima, M., Kawata, S., Abe, M., Kita, Y., Hashimoto, K., Shimizu, T., Watanabe, M., Sakimura, K., & Kano, M. (2010). The endocannabinoid 2-arachidonoylglycerol produced by diacylglycerol lipase alpha mediates retrograde suppression of synaptic transmission. Neuron, 65(3), 320–327. https://doi.org/10.1016/J.NEURON.2010.01.021

Wilson, R. I., & Nicoll, R. A. (2002). *Endocannabinoid Signaling in the Brain*. Retrieved June 18, 2022, from https://www.science.org

Yoshino, H., Miyamae, T., Hansen, G., Zambrowicz, B., Flynn, M., Pedicord, D., Blat, Y., Westphal, R. S., Zaczek, R., Lewis, D. A., & Gonzalez-Burgos, G. (2011). Postsynaptic diacylglycerol lipase mediates retrograde endocannabinoid suppression of inhibition in mouse prefrontal cortex. The Journal of Physiology, 589(Pt 20), 4857–4884. https://doi.org/10.1113/JPHYSIOL.2011.212225

